# Transcriptional bursting explains the noise-versus-mean relationship in mRNA and protein levels

**DOI:** 10.1101/049528

**Authors:** Roy D. Dar, Sydney M. Schaffer, Siddarth S. Dey, Jonathan E. Foley, Abhyudai Singh, Brandon S. Razooky, Adam P. Arkin, David V. Schafer, Michael L. Simpson, Arjun Raj, Leor S. Weinberger

**Affiliations:** Department of Bioengineering, University of Illinois Urbana-Champaign; Carl R. Woese Institute for Genomic Biology, University of Illinois at Urbana-Champaign, Urbana, Illinois, USA; Center for Biophysics and Quantitative Biology, University of Illinois at Urbana-Champaign, Urbana, Illinois, USA; Department of Bioengineering, University of Pennsylvania, Philadelphia, Pennsylvania 19104, USA; Hubrecht Institute-KNAW (Royal Netherlands Academy of Arts and Sciences) and University Medical Center Utrecht, Cancer Genomics Netherlands, Uppsalalaan 8, 3584CT Utrecht, the Netherlands; Gild, Inc., San Francisco, California 94104, USA; Department of Electrical and Computer Engineering, University of Delaware, Newark, Delaware, USA.; Rockefeller University, New York, New York, USA; Department of Bioengineering, University of California, Berkeley, California, USA; Physical Biosciences Division, Lawrence Berkeley National Laboratory, Berkeley, California, USA; Department of Chemical and Biomolecular Engineering, University of California, Berkeley, California, USA.; Department of Cell and Molecular Biology, University of California, Berkeley, California, USA.; Helen Wills Neuroscience Institute, University of California, Berkeley, California, USA; Center for Nanophase Materials Sciences, Oak Ridge National Laboratory, Oak Ridge, Tennessee, USA; Bredesen Center for Interdisciplinary Research and Graduate Education, University of Tennessee, Knoxville, Tennessee, USA; Department of Materials Science and Engineering, University of Tennessee, Knoxville, Tennessee, USA; Gladstone Institute (Virology and Immunology), San Francisco, California, USA; Department of Biochemistry and Biophysics, University of California San Francisco, San Francisco, California, USA.

## Abstract

Recent analysis (Dey et al, 2015), demonstrates that the HIV-1 Long Terminal Repeat (HIV LTR) promoter exhibits a range of possible transcriptional burst sizes and frequencies for any mean-expression level. However, these results have also been interpreted as demonstrating that cell-to-cell expression variability (noise) and mean are uncorrelated, a significant deviation from previous results. Here, we re-examine the available mRNA and protein abundance data for the HIV LTR and find that noise in mRNA and protein expression scales inversely with the mean along analytically predicted transcriptional burst-size manifolds. We then experimentally perturb transcriptional activity to test a prediction of the multiple burst-size model: that increasing burst frequency will cause mRNA noise to decrease along given burst-size lines as mRNA levels increase. The data show that mRNA and protein noise decrease as mean expression increases, supporting the canonical inverse correlation between noise and mean.

**Conflict of Interest:** The authors declare that they have no conflict of interest.

---

A substantial body of literature has reported an inverse relationship between the mean level of gene expression and the variability or ‘noise’ in expression for genes across biological systems ranging from *E. coli* to mammalian cells (Sanchez & Golding, 2013). The noise-mean inverse correlation can be explained by a two-state transcriptional ‘burst’ (a.k.a. ‘random telegraph’) model (Kepler & Elston, 2001; Peccoud & Ycart, 1995) where promoters toggle between active and inactive states with a given ‘burst frequency’ and can generate ≥ one mRNA (the ‘burst size’) during each activation event.

A recent analysis (Dey et al, 2015), demonstrates that the HIV-1 Long Terminal Repeat (HIV LTR) promoter exhibits a range of possible burst sizes and frequencies for any mean-expression level. However, these results have also been interpreted as demonstrating a lack of correlation between noise and mean. Here, we re-examine the available HIV LTR data—and perform a new perturbation experiment—to quantify the noise as mean expression increases. The re-analysis and new data show that expression noise contracts along constrained burst-size manifolds as mean expression increases, supporting the canonical noise-mean correlation.

The theoretical basis for the inverse noise-mean correlation derives from analytical solutions of the two-state model, which can, in the bursting regime, generate ‘manifolds’ or ‘lines’ of constant burst size along which burst frequency varies (Franz et al, 2011; Kepler & Elston, 2001; Singh et al, 2010). For example, for a promoter with low burst frequency (*k*_*off*_ >> *k*_*on*_), increasing the burst frequency increases the mean-expression level but simultaneously decreases noise (typically measured by coefficient of variation, CV or CV^2^). Specifically, the two-state model predicts that the noise reduction from increasing burst frequency scales inversely with the mean. Consequently, on plots of CV versus mean, a specific promoter will be observed to ‘slide’ along a hyperbolic manifold of constant burst size that scales inversely with the mean.

This inverse noise-mean correlation was observed in previous measurements of HIV LTR expression (Dar et al, 2014; Dar et al, 2012; Singh et al, 2010; Skupsky et al, 2010). These measurements of GFP protein expression from the LTR promoter showed that different loci in the human genome generate different burst sizes and frequencies but are constrained along hyperbolic manifolds of constant, integer-valued burst sizes (Fig. 1A), where burst sizes were inferred from quantification of GFP <u>m</u>olecular <u>e</u>quivalents of <u>s</u>olubilized <u>f</u>luorophores (MESF). These hyperbolic manifolds are also present in the clones examined by Dey et al. (2015), when auto-fluorescence is accounted for (Fig. S1).

**Figure 1:**
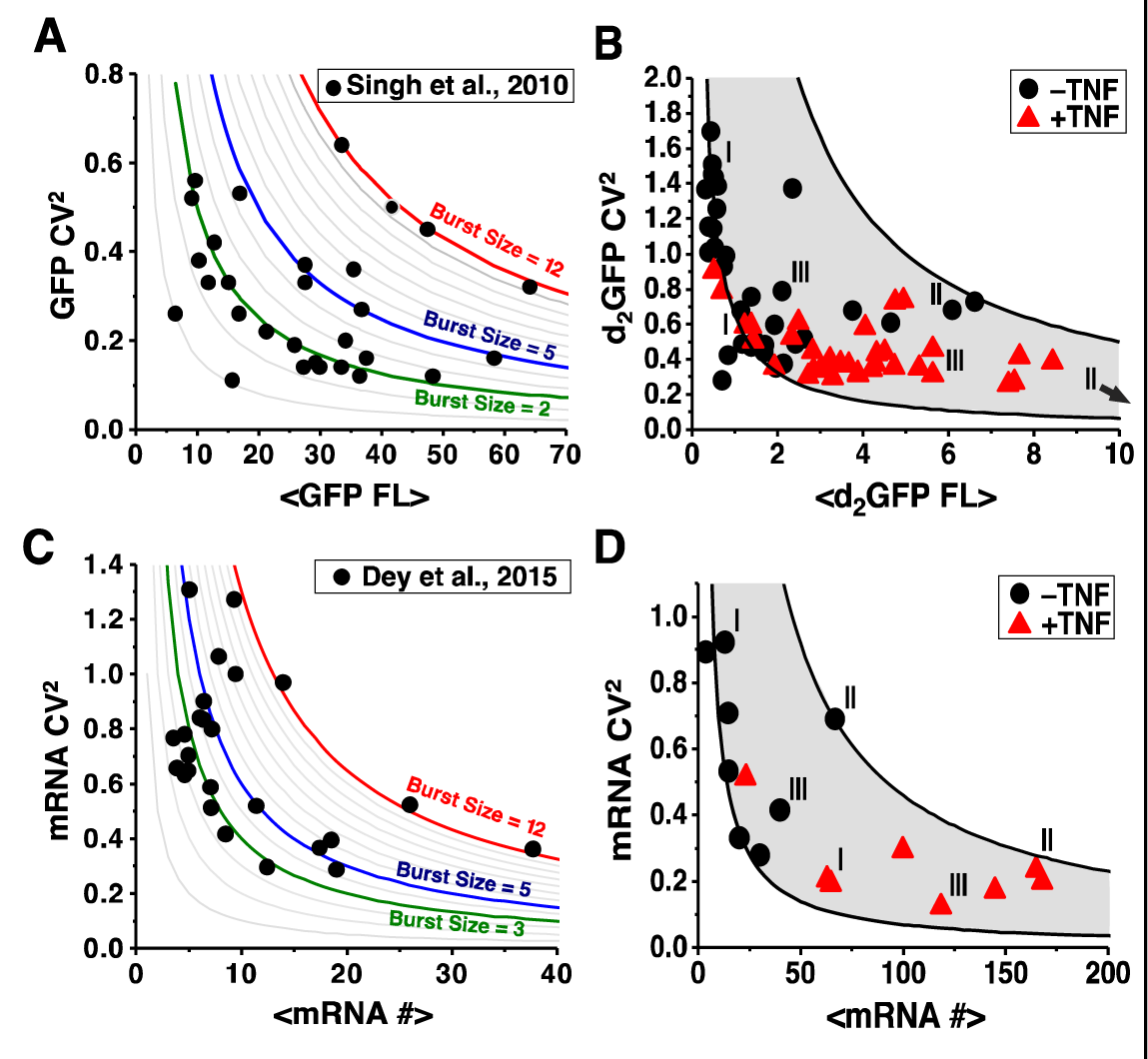
Protein and mRNA noise are inversely correlated with abundance. **(A)** Re-plotting of (Singh et al, 2010) GFP protein measurements for 30 HIV LTR-GFP isoclonal cell populations each with a distinct genomic integration site. Each point represents ~3,000 clonal cells (extrinsic noise filtered out by sub-gating of 50,000) and clones fall along distinct hyperbolic manifolds of transcriptional burst that are analytical solutions to the two-state model where Burst Size = (CV^2^ × <GFP MESF>) / 5,000 - 1 as in (Singh et al, 2010). Grey lines represent burst sizes from 0-12. Color lines are highlighted burst sizes. (**B**) 30 different LTR-d2GFP (2-hr half-life GFP) clonal populations before TNF-α (black) and after 18-hr TNF-α (red) exposure, reproduced from (Dar et al, 2012) where extrinsic noise was filtered out as in A. As predicted from the two-state model, noise is constrained between hyperbolic manifolds of constant burst size (gray). Black lines represent min and max burst size lines fit to dimmest and brightest clones, respectively, before TNF-α exposure. Representative individual clones are labeled as I, II, and III. (**C**) Re-plotting of Dey et al. (2015) smFISH RNA measurements for 23 LTR-GFP isoclones (Burst Size = (CV^2^ × <mRNA #>) showing that clones fall along distinct burst model lines. (**D**) New smFISH analysis of LTR-d_2_GFP mRNA for eight different clones (a subset of isoclones originally reported in (Dar et al, 2012)) before TNF-α (black) and after 18-hr TNF-α (red) exposure. Clones I, II, and III are the same clones as in panel B and black lines calculated as in panel B.

Further measurements validated the prediction that perturbing transcriptional burst frequency confines noise changes between manifolds of constant burst size (Dar et al, 2012; Singh et al, 2010). In vivo, HIV LTR transcription is activated by recruitment of transcription-initiation factors to nuclear factor kappa B (NFκB) sites on the LTR, which is promoted by the inflammatory cytokine <u>T</u>umor <u>N</u>ecrosis <u>F</u>actor α (TNFα). Upon TNFα exposure, LTR expression was found to increase, but in concert with contraction of CV^2^ between constrained manifolds of minimal and maximal burst size (Fig. 1B). As previously reported, there exists an expression-level threshold above which burst size—rather than burst frequency—begins to change (Dar et al, 2012; Skupsky et al, 2010) causing clones to deviate from a single burst-size line at higher expression levels. Nevertheless, CV^2^ is constrained between burst-size manifolds and the inverse noise-mean correlation is preserved (i.e. the extreme upper-right and lower-left regions of CV^2^-vs.-mean space are devoid of data). However, there was potential concern that these measurements were based on protein fluorescence, rather than RNA, where transcriptional burst size could only be inferred from quantitative modeling and MESF.

A powerful method that provides a more direct measure of transcriptional burst size is <u>s</u>ingle-<u>m</u>olecule RNA <u>F</u>luorescence <u>*i*</u>*n* <u>*s*</u>*itu* <u>H</u>ybridization (smFISH), which counts diffraction-limited spots of individual RNA molecules (Raj et al, 2006). Recent analysis of HIV LTR expression comprehensively examined both GFP protein and RNA levels for 23 isoclonal populations (Dey et al, 2015). Here, we analyze this smFISH RNA expression data and find that the isoclonal populations fall along hyperbolic manifolds of constant burst sizes (Fig. 1C). The burst sizes from smFISH range between 2-12 mRNAs with the majority of isoclones exhibiting burst sizes of 2-5 mRNAs, in agreement with the burst-sizes inferred from GFP fluorescence (i.e., burst sizes inferred from GFP range from 2-12, with the majority of isoclones displaying burst sizes of 2-4).

To further test whether expression is constrained to hyperbolic manifolds of constant burst size, we performed smFISH measurements of a subset of eight isoclonal LTR populations before and after TNFα exposure. For all isoclonal populations, TNFα increases the mean number of mRNAs transcribed from the LTR, but at the same time leads to a concomitant contraction of the CV^2^ between constrained manifolds of burst size (Fig. 1D). Overall, these smFISH data support a strong inverse correlation between noise and mean expression.

To summarize, the GFP protein and mRNA analyses are in general concordance both quantitatively, in terms of the burst-size values matching, and qualitatively, in terms of the inverse noise-mean relationship being conserved. While this analysis examines only the HIV LTR promoter, the inverse noise-mean relationship has been observed for a range of promoters (Dar et al, 2012) across different organisms and under varying conditions (Sanchez & Golding, 2013), suggesting that it is a general feature of gene expression. Methodologically, this analysis underscores the reliability of protein-level measurements for quantifying transcriptional parameters (Singh et al, 2012). From an application standpoint, validating the burst-size manifolds lays an important theoretical foundation for explaining how noise enhancers and suppressors synergize with transcriptional activators to modulate fate-selection decisions, such as HIV reactivation from latency (Dar et al, 2014).

## Supplemental Figure S1

**Supplemental Figure S1:**
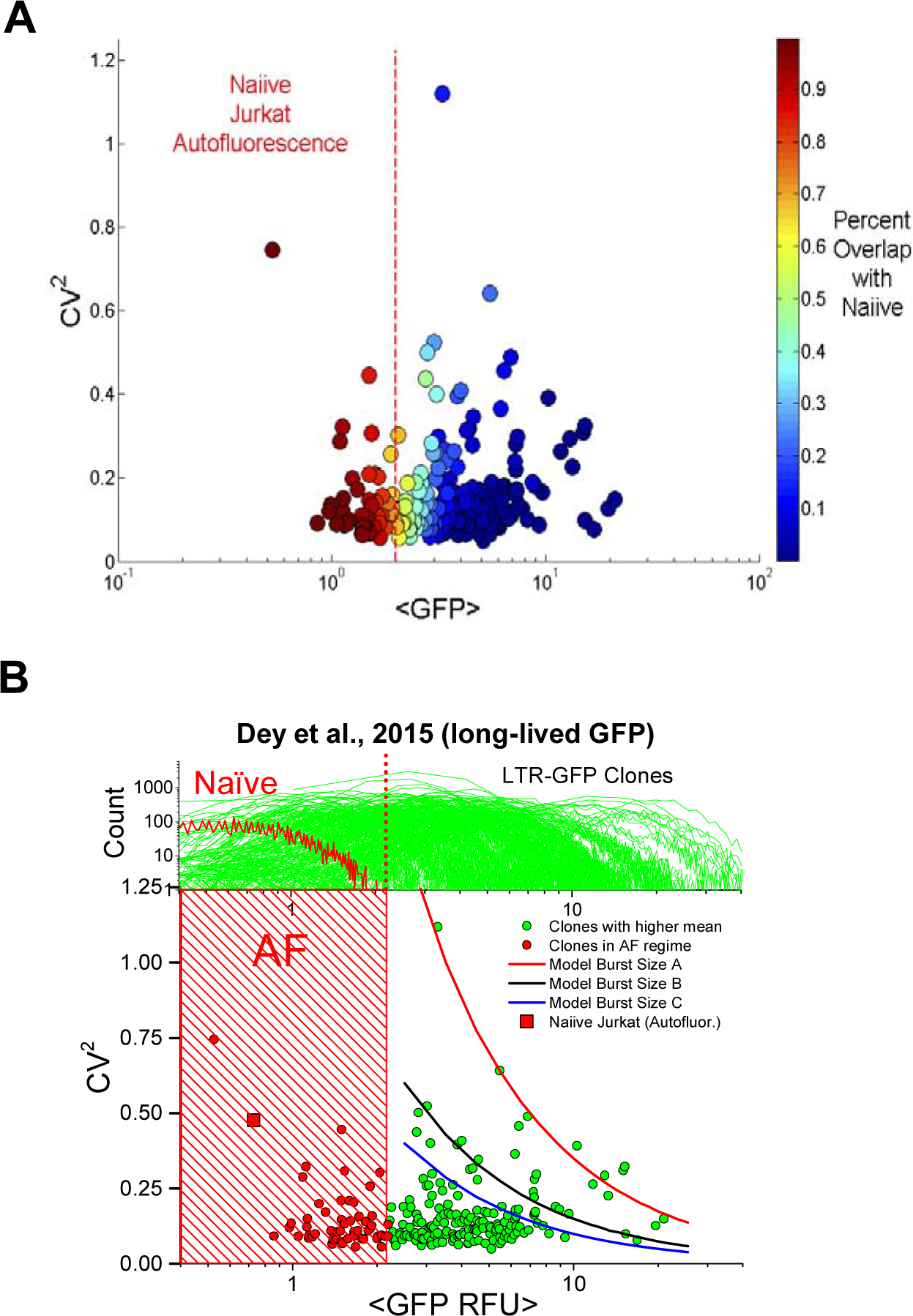
Re-analysis of GFP flow cytometry data. (**A**) Re-plotting of Dey et al. (2015) GFP protein flow cytometry data relative to overlap with autofluorescence regime (200 isoclonal populations are shown). Each point represents ~3,000 isoclonal cells (extrinsic noise filtered out by sub-gating of 50,000). Many clones exhibit significant overlap with the autofluorescence regime. (**B**) The clones farthest from the autofluorescent regime clones fall along hyperbolic manifolds of constant burst size. Lines correspond to CV^2^ = (Q(1 + m)) / <GFP RFU> where m = 1 for blue line, 1.5x for black line, and 3.5x for red line, and Q is a fit constant (burst size cannot be calculated from this data due to lack of absolute quantitation). It is not clear if clones with the lowest CV correspond to lower burst sizes or if the extrinsic noise limit (Taniguchi et al, 2010) dominates at this low CV level.

